# The technique of articular rabbit cartilage roughness measurement and analysis of the effect of PRP in knee osteoarthritis therapy using atomic force microscopy

**DOI:** 10.1101/2020.03.26.009837

**Authors:** Ihnatouski M. Mikhail, Jolanta Pauk, Dmitrij Karev, Borys Karev

**Affiliations:** Yanka Kupala State University of Grodno, Scientific and Research Department, Grodno, Belarus; Bialystok University of Technology, Institute of Biomedical Engineering, Bialystok, Poland; Grodno State Medical University, Department of Traumatology, Orthopedics and Field Surgery, Grodno, Belarus; Grodno City Emergency Hospital, Department of Orthopedic and Traumatology, Grodno, Belarus

**Keywords:** atomic force microscopy, hyaline cartilage, platelet-rich plasma therapy, induced model, osteoarthrosis

## Abstract

Hyaline cartilage undergoes degenerative-dystrophic changes with subsequent involvement of the subchondral bone. The purpose of this study was developing a new AFM-based method to articular rabbit cartilage roughness measurement, followed by an investigation of whether platelet-rich plasma therapy of knee osteoarthritis has a positive impact on mechanical properties of rabbit cartilage. The rabbits were randomly divided into two groups: the control group (N=6) and the patients (N=12). Saline (0.5 ml) and 10% surgical talc solution were injected into the right knee of 12 rabbits to induce osteoarthritis. Six rabbits underwent PRP therapy, while the other six did not receive treatment. The mechanical properties and the submicron surface morphology rabbit hyaline cartilage were investigated using atomic force microscopy (AFM). In the group of specimens worn out by induced osteoarthritis, the maximum arithmetic average of absolute values (*Ra*) change was a 23% increase; the maximum peak height (*Rp*) increased by over 100%, while the mean spacing between local peaks (*S*) increased by 26%, compared to healthy rabbit cartilage (p<0.05). In the group of specimens worn out by induced osteoarthritis and cured with PRP therapy, *Ra* increased by 13%; *Rp* increased by 33%, while *S* decreased by 77%, compared to healthy rabbit cartilage (p<0.05). It was found that the mechanical properties of hyaline cartilage deteriorate under the influence of simulated osteoarthritis. The results of PRP treatment in rabbits may constitute a step forward to further relevant studies involving OA patients.

## Introduction

Osteoarthritis (OA) is a musculoskeletal disease common throughout the world, affecting 10-15% of the population aged 45 years and older [1,2]; over 30 million people in the United States [3] and 302 million worldwide suffer from OA [4]. Currently, osteoarthritis ranks 12^th^ globally among all the causes of disability in the population [4]. The prevalence of osteoarthritis is increasing and it is expected that 78 million adults in the US will have been affected by 2040 [5]. Osteoarthritis is a chronic progressive disease whose causes include imbalances between anabolic and catabolic processes in joint tissues. The risk factors include old age, obesity, and type 2 diabetes mellitus, previous damage, or mechanical overload to the joint. Hyaline cartilage undergoes degenerative-dystrophic changes with subsequent involvement of the subchondral bone. Moreover, chronic synovitis, sclerosis of the joint capsule, meniscal degeneration, and articular and muscular atrophy are also observed. Osteoarthritis is indicated as one of the most common forms of arthritis, with the knee joint the most commonly affected site [6–8].

For a long time, endoprosthesis of the large joints was the leading method of treatment of clinically pronounced degenerative lesions. In the 21^st^ century, autologous chondrocyte transplantation/implantation has become increasingly common in the treatment of large cartilage defects in the knee joint. In cases of chondral or osteochondral lesions, autologous transfer of cartilage-bone cylinders is the only technique to deliver hyaline cartilage to the defect site of weight-bearing joints such as the knee, ankle, hip, or even the elbow [9,10]. In the 20^th^-21^st^ century, orthopaedists’ attention turned to conservative methods of treatment of joint disease, indicating that new modalities should focus on multifunctional treatment and solving a set of biotribological problems, e.g. suppression of inflammation of the synovial tissue and regeneration of the damaged parts of the friction surfaces of cartilage [11,12]. In the 21^st^ century, non-steroidal anti-inflammatory drugs, glucocorticosteroids, hyaluronic acid, as well as unproven alternative therapies, have all been used to stop the progression of inflammation. This has been the standard treatment in patients who failed to respond to non-pharmacological therapy or analgesics [13]. The efficacy of drug therapies, however, is controversial while clinical guidelines remain inconsistent [14]. Despite the availability of experimental clinical data, published in [15,16], no single method of treatment and prevention of osteoarthritis exists. In recent years, the concept of disease-modifying therapies involving the use of chondroprotective drugs that can control the symptoms of osteoarthritis, stop its progression, and cause modification of cartilage structure [17,18] has been investigated. Studies to assess the feasibility of the use of the natural biological environments of the human body are being conducted in parallel with work on synthetic drugs. The main chondroprotective drugs are biological and synthetic equivalents of natural cartilage components. After injection, they remain in the joint cavity only for a short time and on their own cannot maintain the rheological parameters of the joint fluid for long. A disadvantage of the group of drugs in question is their high cost. Current studies involve the use of platelet-rich plasma therapy (PRP) in the treatment of osteoarthritis [19–23]. Some basic, preclinical, and even clinical case studies and trials report the ability of platelet-rich plasma to improve musculoskeletal conditions, including osteoarthritis [24–27]. Nevertheless, there are still concerns regarding its clinical efficiency, mainly due to the heterogeneity of the used methods and the obtained results. The lack of scientific evidence and standardization between different PRP protocols result in the efficacy of platelet-rich plasma therapy remaining unclear.

The pathogenesis, pathophysiology, and the effect of therapeutic intervention in OA treatment requires the development of animal models, which would play a crucial role in the understanding of the disease. Such models are classified in literature on the basis of their aetiology, i.e. as primary osteoarthritis, post-traumatic osteoarthritis, etc. [28]. Both small (mouse, rat, rabbit, and guinea pig) and large animals (dog, goat/sheep, and horse) have been used to develop OA models [29–35]. The most common models include anterior cruciate ligament transection, medial meniscal tear, and meniscectomy. Chemically induced models are mainly used for the purpose of the assessment of the effectiveness of treatment in inflammation. They involve the injection of an inflammatory compound (sodium monoiodoacetate, quinolone, collagenase, etc.) directly into the knee joint with the aim of inducing OA in animals [28]. Studies of histopathological outcomes have been widely published in [26,35]. However, a beneficial impact of intra-articular PRP injection on the mechanical properties and the submicron surface morphology of hyaline cartilage has not been fully recognized.

The current trend in orthopaedics is to develop methods of prevention of diseases of the musculoskeletal system through the development and improvement of invasive and non-invasive diagnostic technologies [36–38]. Radiography, computed tomography, and MRI are used to study various animal models [39–42]. In [43], According to Nieminen, the most common imaging modalities are radiography (N=658) and MRI (N=312), followed by CT (N=182), ultrasound (N=92), and nuclear medicine (N=15); he also shows that imaging studies play an increasingly important role in osteoarthritis research. However, studies of mechanical changes and submicron surface morphology of hyaline cartilage in the knee joint require high-resolution force scanning techniques [28, 44–46]. Electron microscopy reveals ultrastructural details at a molecular resolution, its disadvantage being the complexity and lengthy sample preparation procedures. Moreover, neither light nor electron microscopy can measure the mechanical properties of cartilage directly. In contrast, atomic force microscopy (AFM) enables simultaneous imaging and measurement of stiffness on the micrometer-nanometer scale in native samples, thus helping to elucidate the structure and the mechanical properties of articular cartilage. Hence, the purpose of this study was developing a new AFM-based method to articular rabbit cartilage roughness measurement, followed by an investigation of whether platelet-rich plasma therapy of knee osteoarthritis has a positive impact on mechanical properties of rabbit cartilage.

## Methods

The experiments on animals were performed in accordance with internationally accredited guidelines, and had been approved by the Ministry of Health of the Republic of Belarus (No. 3, 2017).

### Experimental animals and PRP preparation

Eighteen rabbits (chinchilla) from the Faculty of Veterinary Medicine at Grodno State Agrarian University, aged five months and weighing from 2,400 to 3,500 g, were included in the study. They had free access to food and water during the experiments. The inclusion criteria were good appetite and activity, a shiny even coat, a clean nose and eyes, a body temperature in the range 38.5–39.5 degrees, a heart rate ranging from 120 to 160 BPM, and a respiratory rate ranging from 50 to 60 movements per minute. The exclusion criteria included lack of appetite, intestinal disease, inactivity, or suppressed behaviour. Before the start of the experiments, the rabbits were randomly divided into two groups: the control group (N=6) and the patients (N=12). 9 ml of blood was drawn from the marginal vein of the ear of each of the rabbits and mixed with 0.5 ml of 0.0775 mol/L sodium citrate acting as an anticoagulant. The blood was centrifuged at 1,200 rpm (160 g) for 20 minutes to separate the plasma containing platelets from red cells. The plasma was drawn from the top layer and centrifuged for an additional 15 minutes at 2,000 rpm (450 g) to separate the platelets at a temperature of +18 to +22°. After centrifugation, 0.6-0.7 ml of PRP was drawn from a plasma layer rich in platelets using a syringe and an injection needle. The obtained PRP contained growth factors, whose concentration was about 5 times higher compared to the physiological concentration, which explains their effect on the regeneration process [47].

### Induced animal model of osteoarthritis

The rabbits were immobilized in a special chamber and were anesthetized by intramuscular administration of 10 mg/kg of Xylazine (Ksilanit^®^, Registration Certificate 44-3-2.15-2588NoΠBP-3-8.9/02531 NITA-FARM, Russia) and 50 mg/kg of ketamine (Ketamine^®^, Registration Certificate No. PN000298 / 01, Moscow endocrine plant, Russia) which kept the animal sedated for a longer time with minimal pain. Xylazine coadministered with ketamine was a safe anesthetic adjunct to induce short periods of surgical anesthesia. In [48], it was proved that the combination of both Xylazine and ketamine cause rabbits’ muscle relaxation; rabbits have a smoother emergence from anesthesia. Saline (0.5 ml) and 10% surgical talc (Talcum Pharmaceutical^®^, “SNABPISHEPROM” LLC, Belarus) solution were injected into the right knee under the patella to induce osteoarthritis in 12 rabbits. Talc is a clay mineral composed of hydrated magnesium silicate. We used surgical talcum powder as a trademark to distinguish it from E553b and other types of talcum powder. Six rabbits underwent PRP therapy, while the other six did not receive PRP treatment. Four injections 0,5 ml PRP were administered in 7-day intervals. The animals were euthanized on the 28^th^ day injection. Rabbit articular cartilage was harvested from the femoral condyles by cutting specimens off the underlying bone with a sharp razor blade, yielding ~5 mm × 5 mm pieces, ~2 mm thick.

### Analysis of submicron surface morphology in rabbits

The rabbit specimens were fixed on a rigid substrate and stuck to microscope slides. Rabbit cartage specimens were selected based on an initial visual assessment of surface degradation at optical magnifications of 100x, 200x, and 500x. The images were obtained in reflected light using a ©Micro 200T-01 optical microscope. Tests of submicron surface morphology of rabbit specimens were performed using AFM NT-206 (©MicroTestMachines, Belarus) in the static scanning mode. A CSC38 MikroMasch^®^ silicon probe was used; the resulting tip radius was less than 35 nm; the full tip cone angle was 40°; the total tip height was 12 to 18 μm; the probe material was n-type silicon; a type A cantilever was used. The resonance frequency of the cantilever ranged from 8 to 32 kHz; the force constant was 0.01 to 0.36 N/m; length 250±5 μm; width 32.5±3 μm; thickness 1.0±0.5 μm. The areas of specimens without surface waviness or high strain amplitudes were found prior to scanning or indenting by means of an optical microscope built into the AFM. The scanning results were assessed at medium magnification with a scan area of *Ar*=9×9 μm^2^. The degree of surface wear was determined by measuring the roughness parameters. The SurfaceExplorer (©MicroTestMachines) software and nano images (©Mikhail Ihnatouski) were used to visualize the experimental data and to measure the roughness parameters.

Specimens obtained after division into the above two groups were used for further studies of submicron surface morphology and the mechanical properties. Measurements of the roughness parameters of cartilage surfaces were performed, including the arithmetic average of absolute values (*Ra*), the maximum peak height (*Rp*), and the mean spacing between local peaks (*S*). The roughness parameters were measured at 5 points of each of the 25 specimens for three different values of *Ar*.

### Analysis of the mechanical properties of rabbit cartilage

The radius of AFM tip and stiffness of cantilevers were calibrated. Force spectroscopy, i.e. AFM, was used to measure the mechanical properties of specimens of the rabbit cartilage. The measured quantities were the bend of the console (*Z_defl_*) and the displacement of the console along the vertical axis (*Z_pos_*). The penetration of the probe into cartilage (Fig.1) is presented as (2):

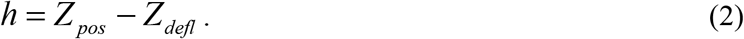

**Fig 1.**
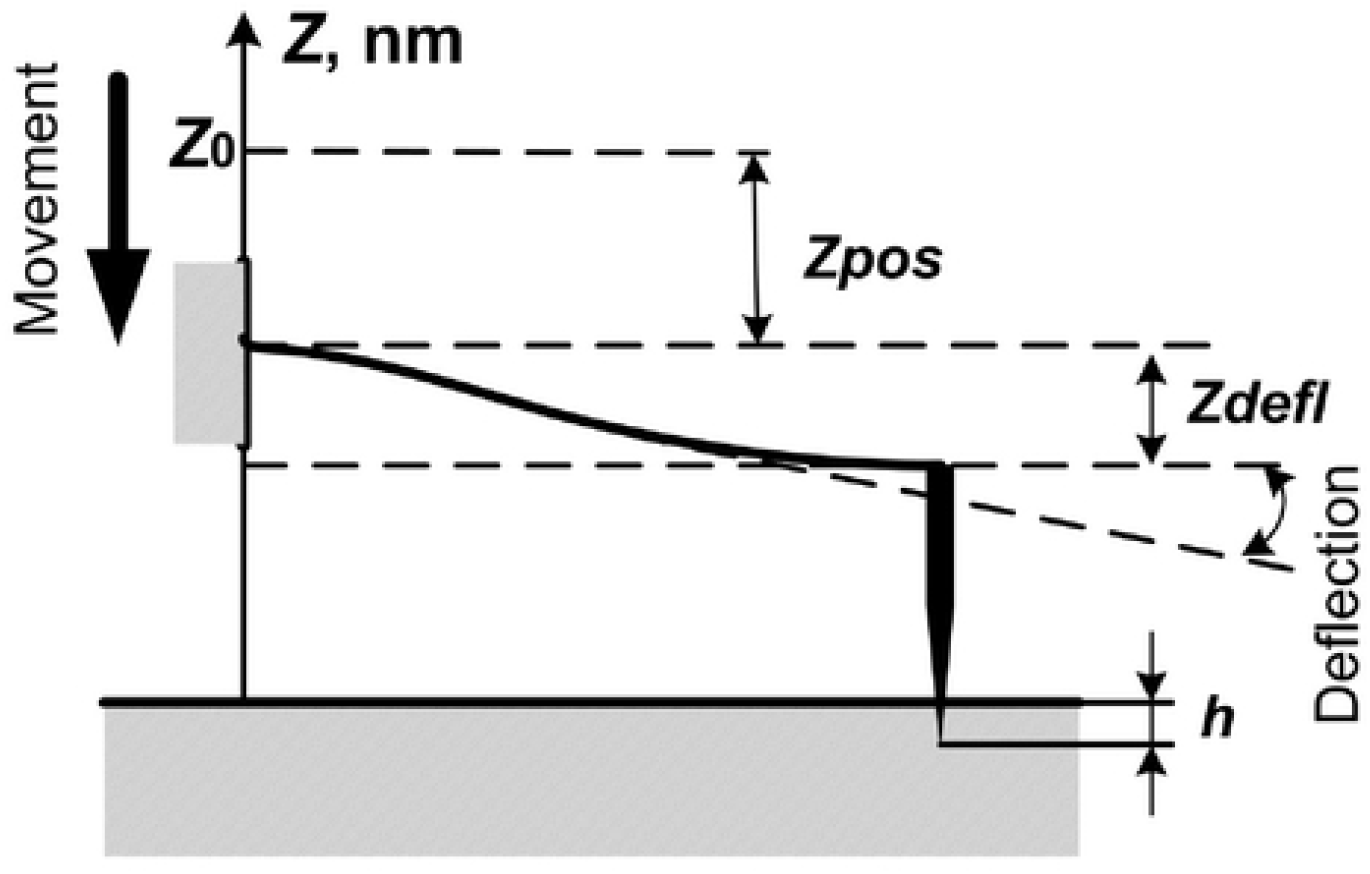
Scheme of AFM indentation.

This can be used for calculating the Young’s modulus (*E*) at a point on the surface [49]:

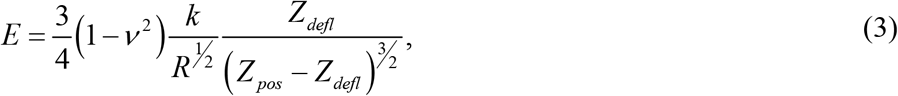

where: *v*=0.5 is the Poisson’s ratio of the cartilage, *k*=0.08 N/m is the stiffness of the cantilever of CSC 38, and *R*=30 nm is the radius of the needle-point of the tip of CSC 38. Thus, the results of the measurements were the relationship between the Young’s modulus and the depth of penetration into the surface (*E*(*h*)).

The surface of each sample was indented using AFM in the course of three measurements, through the introduction of an indenter at a selected point on the surface. The obtained values of Young’s modulus were averaged.

### Statistical analysis

Statistical analyses were performed using the Statistica software (StatSoft 13.1, Cracow, Poland). The data concerning the roughness parameters were checked for normality using the Kolmogorov-Smirnov test. The comparison of the different experimental groups was performed using the Mann-Whitney test. P-value<0.05 was considered as statistically significant. All data are presented as a mean ± standard deviation.

## Results

### AFM monitoring of rabbit cartilage after PRP therapy

Within several minutes following the injection, all the animals were conscious and started to move. None of the rabbits showed any sign of a knee infection or swelling, or died during the test. From the first day, the rabbits developed synovitis, manifested as an increase in the local temperature of the joint. Changes in the morphology of rabbit cartilage were investigated. AFM images of the worn surfaces of healthy rabbit cartilage, after the induction of osteoarthritis and PRP therapy are shown in fig. 2. Damage and gaps were found on the cartilage tissue worn out by Induced Model Osteoarthrosis (IMO).

**Fig 2.**
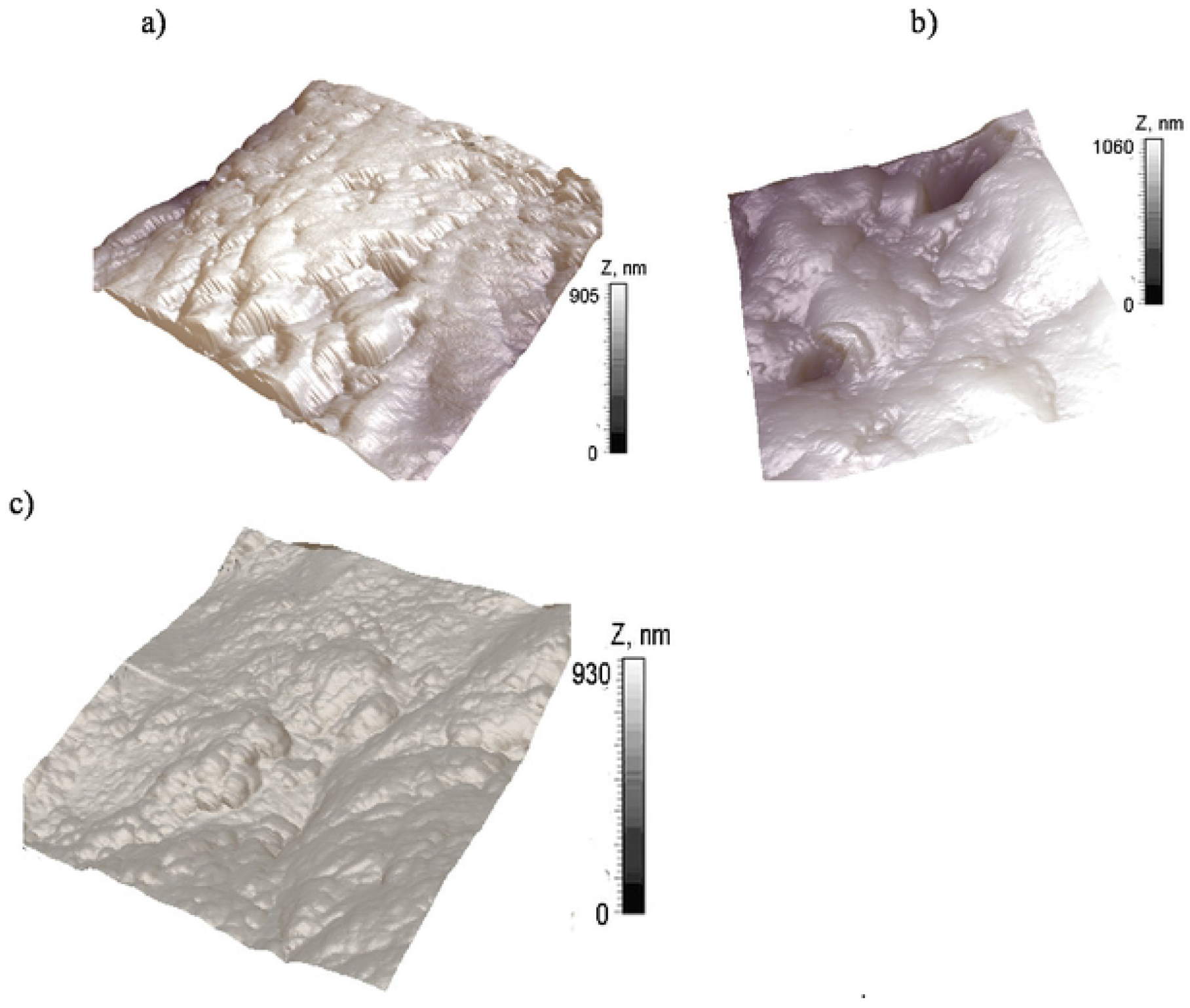
AFM images of rabbit cartilage surfaces 9×9 μm^2^: a) Healthy rabbit cartilage, b) Induced Model of Osteoarthrosis, c) after PRP therapy.

Damage and gaps were found on the cartilage tissue worn out by IMO. The destruction sites were also found on cartilage after PRP therapy. However, the size of the lesions decreased after PRP therapy.

### Comparison of the roughness parameters and the mechanical properties between rabbits groups

The study included 6 healthy rabbits, 6 rabbits whose cartilage was worn out by IMO, and 6 rabbits that received PRP therapy. 30 specimens were obtained from each of the groups (90 specimens in total). The roughness parameters were measured at 5 points of each of the specimens at *Ar*=9×9 μm^2^ (450 measurements in total). The measurement results for a single specimen were averaged. The roughness parameters of cartilage are presented as arithmetic mean (mean) and standard deviation (std), tab. 1.

**Table 1.** Mean (SD) and range of roughness parameters in three groups

| Specimen (N=90) | $R_a$ [nm] | $R_p$ [nm] | $S$ [nm] |
| --- | --- | --- | --- |
| Healthy (N=30) | 64 (3) | 485 (27) | 2005 (92) |
| Worn out by IMO (N=30) | 73 (6)* | 1051 (122) * | 2241 (134) * |
| Worn out after PRP therapy (N=30) | 69 (4) * | 617 (57) * | 1502 (49) * |

\*p<0.05

In the group of specimens worn out by induced osteoarthritis, the maximum *Ra* change was a 23% increase; *Rp* increased by over 100%, while *S* increased by 26%, compared to healthy rabbit cartilage (p<0.05). Thus, the wear caused by artificial osteoarthritis differs from the submicron surface morphology that appears during the natural genesis and course of osteoarthritis. This type of wear can be explained by mechanical damage caused by injections of surgical talc and the short duration of illness (weeks). In the group of specimens worn out by induced osteoarthritis and cured with PRP therapy, *Ra* increased by 13%; *Rp* increased by 33%, while *S* decreased by 77%, compared to healthy rabbit cartilage (p<0.05). The same values amounted to a *Ra* decrease of 8%, *Rp* decrease of 43%, and *S* decrease of 39%, compared to the group of specimens worn out by induced osteoarthritis without PRP therapy (p<0.05). The altitude parameters decreased below the initial value, as did the mean spacing between local peaks, which can be regarded as an increase in the density of small surface irregularities and the smoothing out of large ones. Thus, a restoration of the surface morphology as a result of therapy was recorded. Diagrams showing the relationship between the Young’s modulus and the indentation depth are shown in Fig. 3.

**Fig 3.**
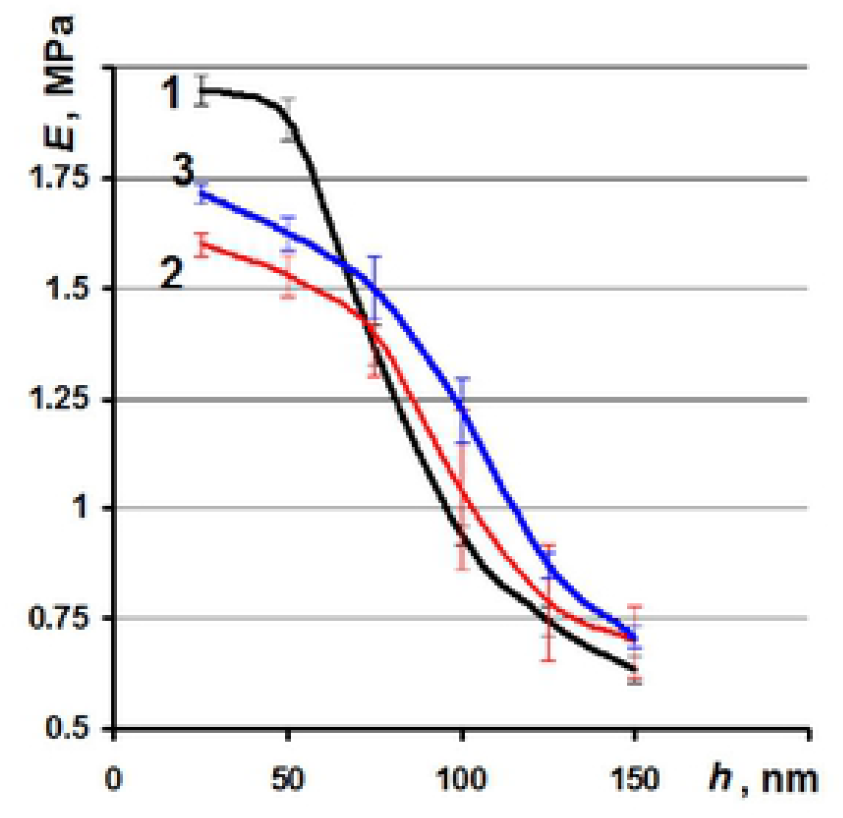
Relationship between the Young’s modulus and indentation depth: line 1 – healthy; line 2 – worn out by IMO; line 3 – worn after PRP therapy.

The values of the Young’s modulus of healthy cartilage (lines 1) ranged from 1.95 MPa on the surface to 0.64 MPa at a depth of 150 nm. The range of the Young’s modulus of the rabbit cartilage worn out by IMO (line 2) was from 1.6 to 0.7 MPa; the range of the Young’s modulus after PRP therapy was from 1.72 to 0.7 MPa. Thus, it was found that the mechanical properties of hyaline cartilage deteriorate under the influence of simulated osteoarthritis. The Young’s modulus of the surface layers decreased by 18% (p<0.05). The AFM showed that the Young’s modulus increased by 9% (p<0.05) as a result of PRP therapy, which means that the mechanical properties of cartilage were partially recovered. The ratio between the Young’s modulus and the indentation depth of healthy and worn rabbit cartilage is steadily decreasing.

## Discussion

Animal models play a significant role in understanding OA and the effects of therapeutic interventions. Authors use both small (mouse, rat, rabbit, and guinea pig) and large animals (dog, goat/sheep, and horse) to develop OA models [29–35]. In this study, an induced model of osteoarthritis, causing the initial stages of primary osteoarthritis in rabbits, was tested. In the opinion of the authors, the rabbit model is the one most adequate for the purpose of this study due to the fact that rabbits have large enough joints and are of a size appropriate for simple surgical procedures and handling of specimens with a cartilage thickness of 0.25 mm-0.75 mm [65]. Rabbits are very often used in the assessment of cartilage repair [22, 32, 54, 60, 61] because the size of chondrocytes in human and rabbit articular cartilage do not differ significantly from each other [62].

The PRP has been recognized as a biological treatment aimed at the regeneration of impaired tissues [50–58]. It is defined as a portion of the plasma fraction of autologous blood with a platelet concentration above the baseline (before centrifugation). In this study, the aim of the proposed modelling was to assess the effect of the treatment agent (the biological environment of the platelet-rich plasma modified in vivo and in vitro) on the knee joint. PRP therapy of induced osteoarthritis was developed and tested. Growth factors play a crucial role in bone tissue regeneration, while PRP therapy results in the highest concentration of platelet levels in blood. When compared to the total blood concentration, the platelet concentration for PRP obtained in this study was about 5.0 times higher than the baseline, which was sufficient to improve bone regeneration. Some authors propose concentrations 4-6 times higher than the baseline [52,53]. Saline (0.5 ml) and 10% surgical talc solution were injected into the right knee under the patella to induce osteoarthritis as the knee is one of the most common sites of musculoskeletal pathologies in OA. In some studies, sodium, quinolone, or collagenase were delivered directly into the knee joint with the aim of inducing OA in animals [28]. The results of this study show that PRP injection into the right knee under the patella could improve the mechanical properties and the submicron surface morphology of hyaline cartilage in rabbits. The positive effect of PRP therapy was revealed in AFM images in the group of specimens worn out by the induced osteoarthritis and cured with PRP therapy: *Ra* increased by 13%, *Rp* increased by 33%, while *S* decreased by 77% compared to healthy rabbit cartilage (p<0.05). Moreover, *Ra* decreased by 8%, *Rp* decreased by 43%, and *S* decreased by 39% compared to the group of specimens worn out by induced osteoarthritis without PRP therapy.

The study shows that the Young’s modulus of cartilage worn out by induced osteoarthritis ranged from 1.6 to 0.7 MPa, while the values of the Young’s modulus after PRP therapy ranger from 1.72 to 0.7 MPa. The Young’s modulus increased by 9% as a result of PRP therapy, which means that the mechanical properties of cartilage partially recovered. Indentation performed using an AFM silicon probe clearly revealed a decrease in the Young’s modulus of the upper layers of hyaline cartilage (up to 100 nm). These findings are consistent with previous studies, which show a positive effect of PRP on cartilage defects and tissue regeneration [50,51,52,55]. It was proved that intra-articular PRP injection could potentially serve as the source of joint lubrication [53]. Intra-articular PRP injections significantly suppress the progression of OA in the rabbit model [54]. In [64,65], intra-articular PRP injection produced positive results of the use of PRP to reduce inflammation in the guinea pig model of OA of the knee. Moreover, PRP was proven useful in attenuating arthritic changes in the porcine model of antigen-induced arthritis [66]. Several studies showed [67,68] that articular injection of PRP could reverse or protect against pathological events in cartilage and synovium based on the mouse model of glucocorticoid-induced femoral head osteonecrosis. Several studies reported successful treatment with PRP in terms of function, with decreased cartilage damage [57,58]. Some authors believe, however, that PRP has short-term effects in patients with articular degeneration and the improvement diminishes during long-term follow-up, which limits the clinical application of PRP as a first-line OA treatment [59,63]. To the best of the authors’ knowledge, this is the first study to assess the response to treatment consisting in intra-articular PRP injection on the mechanical properties and the submicron surface morphology of hyaline cartilage.

Radiography, currently the gold standard for OA imaging, has its limitations in the area of the resolution required for cartilage assessment. The limitations of radiographs are dealt with by using different techniques, such as MRI or ultrasound, which enables in vivo visualization of the quality of cartilage and bone structure as well as all articular and peri-articular tissue. In this study, atomic force microscopy (AFM) was used to test the mechanical properties and the submicron surface morphology of hyaline cartilage in rabbits. The roughness parameters of hyaline cartilage surface: the absolute value (Ra), the maximum peak height (Rp), and the mean spacing between local peaks (S) were measured firstly for healthy rabbits, then worn out by IMO, and finally worn out after PRP therapy. AFM measurements were performed using size of scan window: Ar: 9×9 μm2.

The limitations of this study included the sample size. Moreover, the influence of the physical performance of the rabbits on cartilage regeneration was not considered. Further studies will aim at evaluating the effect of PRP at different platelet concentrations. In the future, the method of activated blood plasma preparation and the protocol for its use will be changed. The use of a liquid cell for AFM measurements is also being contemplated. A liquid cell eliminates surface tension and makes it possible to perform cartilage measurements immediately after obtaining the material.

## Conclusions

The proposed AFM method enables measurement of the impact of the mechanical properties of hyaline cartilage on the stage of wear of its surface. The results obtained are stable and well reflect the direction and magnitude of changes in the properties of cartilage. This study reveals that the damage and gaps of the mechanical properties and the submicron surface morphology of hyaline cartilage were smaller in the group treated with PRP compared to the group that did not receive PRP treatment. This suggests that PRP injection demonstrates the effect of cartilage regeneration in the rabbit knee with induced OA. Regardless of the limitations of this study, the proposed model is useful in improving the understanding of OA. The results of PRP treatment in rabbits may constitute a step forward to further relevant studies involving OA patients.

## Acknowledgments

This work was co-financed by Ministry of Science and Higher Education of Poland within the framework of projects (WZ/WM/12/2019).

## Authors contribution

Funding acquisition: MI JP DK. Methodology: MI JP DK BK. Data curation: DK BK. Investigation: MI BK. Formal analysis: MI JP DK BK. Writing – original draft: MI JP.

